# Ocean Acidification Amplifies the Olfactory Response to 2-Phenylethylamine: Altered Cue Reception as a Mechanistic Pathway?

**DOI:** 10.1101/2020.10.08.329870

**Authors:** Paula Schirrmacher, Christina C. Roggatz, David M. Benoit, Jörg D. Hardege

## Abstract

With carbon dioxide (CO_2_) levels rising dramatically, climate change threatens marine environments. Due to increasing CO_2_ concentrations in the ocean, pH levels are expected to drop by 0.4 units by the end of the century. There is an urgent need to understand the impact of ocean acidification on chemical-ecological processes. To date, the extent and mechanisms by which the decreasing ocean pH influences chemical communication are unclear. Combining behaviour assays with computational chemistry, we explore the function of the predator related cue 2-phenylethylamine (PEA) for hermit crabs (*Pagurus bernhardus*) in current and end-of-the-century oceanic pH. We demonstrate that this dietary predator cue for mammals and sea lampreys is an attractant for hermit crabs. Furthermore, we show that the potency of the cue increases at pH levels expected for the year 2100. In order to explain this increased potency, we assess changes to PEA’s conformational and charge-related properties as one potential mechanistic pathway. Using quantum chemical calculations validated by NMR spectroscopy, we characterise the different protonation states of PEA in water. We show how protonation of PEA could affect receptor-ligand binding, using a possible model receptor for PEA (human TAAR1). Investigating potential mechanisms of pH dependent effects on olfactory perception of PEA and the respective behavioural response, our study advances the understanding of how ocean acidification interferes with the sense of smell and thereby might impact essential ecological interactions in marine ecosystems.

## Introduction

Chemical signalling mediates behaviour, development and physiology in many marine ecosystems (Hay 2009). However, the chemical marine environment is changing rapidly due to ocean acidification (Doney et al. 2009). The continuing uptake of atmospheric carbon dioxide (CO_2_) into the ocean changes the seawater carbonate chemistry and reduces the pH. As a consequence, global average ocean pH has already decreased by more than 0.1 since pre-industrial times to pH 8.1 and is predicted to drop further to pH 7.7 by the end of the century (Bopp et al. 2013; IPCC 2014).

This is of highest concern for marine animal behaviour (Clements and Hunt 2015). Although Clements and Hunt (2015) report primarily negative impacts of ocean acidification on the behaviour of marine organisms, the direction and magnitude of change depend on the species, ecosystem and type of behaviour, with some even improving in efficiency under ocean acidification conditions.

It is particularly important to advance our understanding of the mechanisms behind altered predator-prey interactions under ocean acidification, as they are crucial for survival, social interactions and learning (Ferrari et al. 2010; Draper and Weissburg 2019). Predator kairomones and dietary cues shape trophic cascades, structure communities and mediate food webs (Hahn et al. 2019; Cohen and Forward Jr 2003; Poulin et al. 2018). However, only few predator cues are known in aquatic environments. To expand our knowledge, we draw an example from the terrestrial environment and choose to work with 2-phenylethylamine (PEA), a known dietary predator odour that is detected in the urine of most mammals (Ferrero et al. 2011). In aquatic systems, sea lampreys are known to avoid the smell of PEA (Imre et al. 2014; Di Rocco et al. 2016). Although PEA has been suggested for pest control of sea lampreys (Siefkes 2017), little is known about its role in aquatic environments. Many freshwater and marine algae are known to produce neurotransmitter-like compounds such as PEA (Van Alstyne et al. 2019). In marine environments, PEA has been reported from brown and red macroalgae in Germany and Turkey (Steiner and Hartmann 1968; Percot et al. 2009). Thereby, PEA has been hypothesised to function as a feeding deterrent (Smith 1977). The presumed function of PEA as a deterrent in aquatic chemical communication makes it an ideal infochemical to study the effects of ocean acidification on predator-prey relationships.

In the present study, we work with marine hermit crabs (*Pagurus bernhardus*) to determine the role of the predator-associated cue PEA in present and future pH conditions. We hypothesise that the response to PEA is pH dependent, with an effect observable within the range of ocean acidification. We then investigate the relevance of different possible pathways by which pH can interfere with the observed behaviour. Mechanistically, info-disruption associated with ocean acidification has been linked to GABA receptor functioning in fish: the internal compensation for elevated CO_2_ conditions leads to altered brain ion gradients, interfering with neurotransmitter function (Nilsson et al. 2012; Williams et al. 2019). Electrophysiological and transciptomic measurements also revealed an impaired olfactory system in elevated CO_2_ conditions (Porteus et al. 2018). Furthermore, protonation through pH variation associated with climate change scenarios can change the structure and function of signaling cues and thereby affect olfactory perception (Roggatz et al. 2016; Brown et al. 2002). In this study, we consider four mechanisms by which the decreased pH can lead to an altered hermit crab behaviour (Fig. 1). We discuss the effect of a decreased pH on the signal source (pathway 1), quantify the role of the direct effects of pH on the signalling molecule and its interaction with the receptor (pathway 2 & 3) and discuss a potential interference of ocean acidification with signal transduction in hermit crabs (pathway 4).

**Figure 1:**
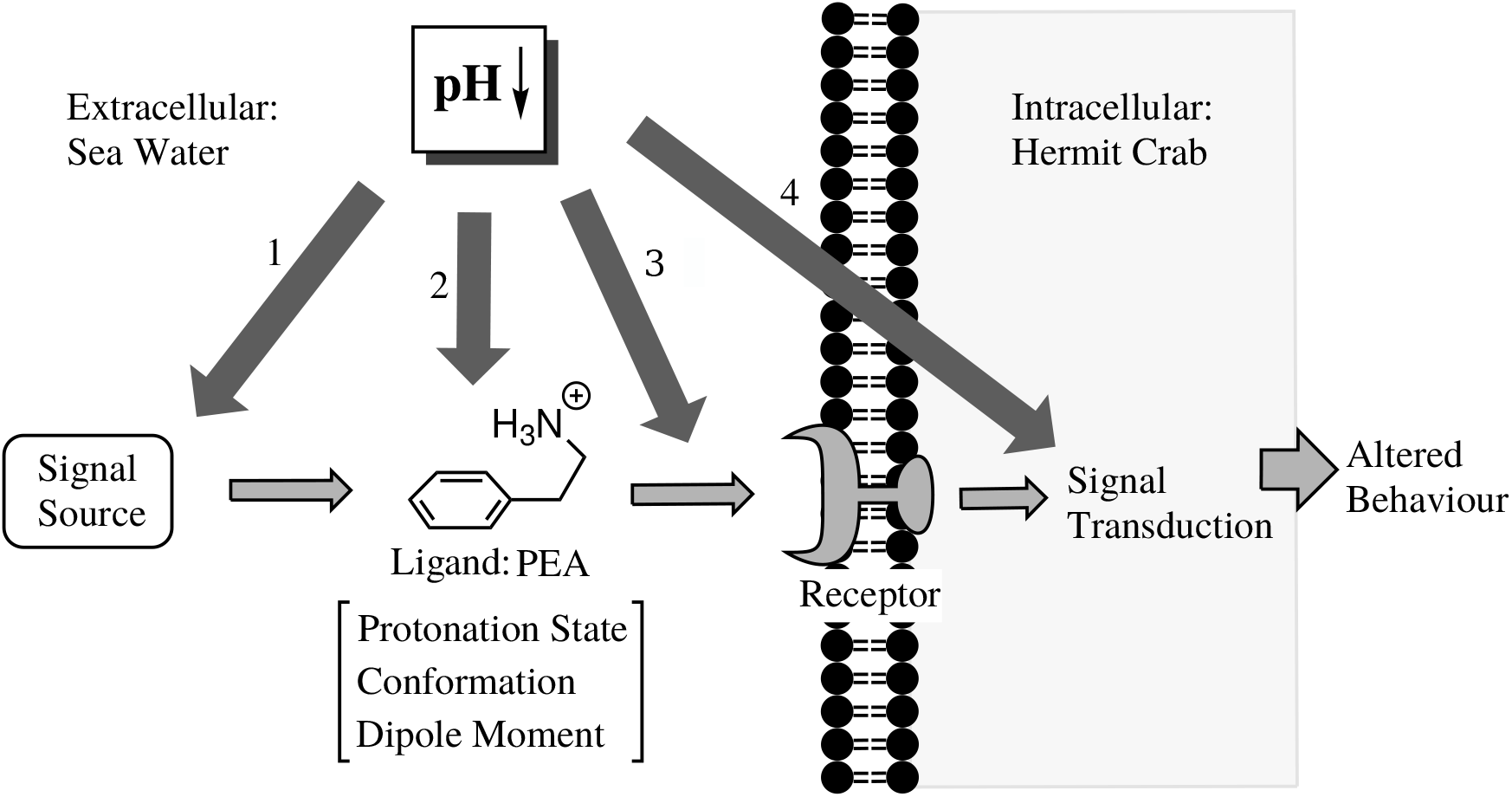
Visualisation of the possible mechanisms by which decreased pH can result in an alteration in hermit crab response to PEA. The pathway of signal transmission from source to response is shown with light grey arrows. The potential mechanisms of a decreased pH interfering with this pathway are shown in dark gray and numbered. Thereby the decreased pH can affect the signal source (1), the signalling cue (2), the receptor or its interaction with the ligand (3) and the signal transduction (4). In this study, the hypothesised scenarios are pathway 2 & 3, whereby the decreased pH alters crucial characteristics of the signalling molecule and subsequently its interaction with the receptor. This ultimately affects the behavioural response.

To advance our understanding of the underlying mechanisms of changing hermit crab response to PEA at decreased pH, we use a range of quantum chemical methods to model the conformation, charge distribution and dipole moment of PEA in different pH conditions (see pathway 2 in Fig. 1).

To date, the conformation of PEA (see Fig. 2) has mostly been studied in gas phase in the uncharged state: rotational and infrared spectra found a strong preference for folded *(gauche)* conformations (Godfrey et al. 1995; López et al. 2007). Thereby, the interaction of the amino group with the *π*-system contributes largely to the stabilisation (Chiavarino et al. 2014; Bouchet et al. 2015). A direct interaction of PEA and one water molecule was found when 1:1 PEA-water clusters were identified in gas phase (Dickinson et al. 1998; Hockridge and Robertson 1999). Further studies applied infrared and rotational spectroscopy paired with computational chemistry to confirm a *gauche* 1:1 PEA-water complex (Melandri et al. 2010; Bouchet et al. 2016). However, recently, the hydration properties of neutral and protonated PEA (PEAH^+^) were also assessed in molecular dynamics simulations revealing a potential conformational preference for *gauche* PEA-water clusters for both protonation states (Ristić et al. 2019). Although PEA is found in aquatic environments, to the best of our knowledge, no study has yet addressed solvation effects on the potential energy curve of both protonation states of PEA. We hypothesise that both the conformation and charge distribution of PEAH^+^ differ from its neutral state and anticipate solvation plays a significant role.

**Figure 2:**
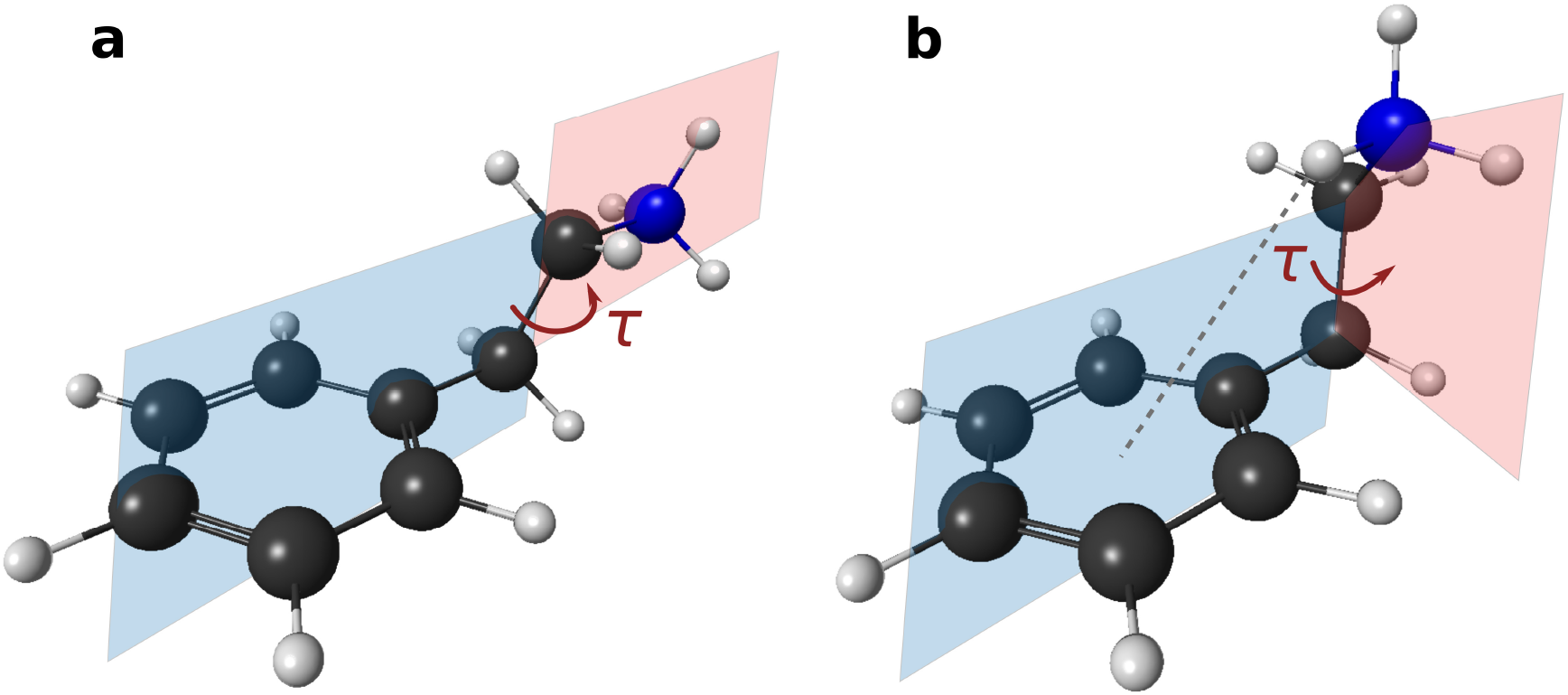
Conformations of protonated PEA (PEAH^+^) with torsion angle *τ* of the amino side chain in dark red. The *anti* conformation (a) correspond to an extended geometry with *τ* ≈ 180°. The torsion angle of the folded conformation (*gauche*, b) is *τ* ≈ ±60° and leads to a weak *π*-hydrogen bond (dotted line).

To relate the computational method to the reality of PEA in water, we compare its calculated structure to NMR spectroscopic measurements. This allows us to validate the computational results.

We also explore the effect of ligand protonation on the receptor binding using quantum chemical calculations. To provide a proof a principle, we study the interaction of PEA with the binding pocket of the human trace amine-associated receptor 1 (TAAR1), whose primary endogenous ligand is PEA. Electrostatic interactions are essential for receptor-ligand interactions (Leckband et al. 1992) and protonation of the chemical signal can affect its electronic properties (Radić et al. 1997). We hypothesise therefore, that the protonation of PEA substantially alters its receptor binding properties. To the best of our knowledge, this is the first study to explore the effect of ocean acidification on receptor-ligand interactions.

This study evaluates the effect of PEA on a marine crustacean at current average pH and end-of-the-century levels, linking behavioural assays with computational chemistry. To determine the functionality of the cue we start by exploring behavioural reactions of hermit crabs in different pH conditions. These lead us to examine the properties of the different protonation states of PEA in water using quantum chemical calculations. The computational findings are then compared with measurements of PEA in water using NMR spectroscopy. Ultimately, we assess the effect of protonation on the interaction of PEA with the potential receptor model TAAR1 and draw conclusions on the possible mechanisms by which a decreasing ocean pH can affect the behavioural response to a chemical signal.

## Materials and Methods

### Hermit Crab Collection and Culture

*P. bernhardus* were collected by hand from the rocky intertidal shore near Scarborough (54°25’19.6”N 0°31’43.6”W), UK, in November 2018. At the aquaria facilities of the University of Hull, the hermit crabs were kept at pH 8.1 ± 0.1 and acclimatised to a twice weekly feeding rhythm with cooked blue mussels and kept at an average temperature of 15.8 ± 0.2 °C and 35.9 ± 0.2 PSU.

### Behavioural Assay to Determine Cue Functionality

To determine the reaction of hermit crabs to PEA, the animals (n=20 per pH condition) were starved for 5-7 days and randomly allocated to be tested in pH condition 7.7 and 8.1. The light was dimmed to reduce the impact of visual stimuli. After being acclimatised to the pH of the new environment for up to 2 minutes in a separate tank, each individual was tested for its reaction to PEA (2-phenylethylamine hydrochloride, Sigma-Aldrich, 98%). Thereby, three concentrations were tested subsequently with up to 2 minute acclimation time in a seperate tank between the assays. As negative control, the undisturbed movement pattern of each individual in their allocated pH condition and tank was observed in an experiment without chemical cues. This experiment helps to identify a potential bias of movement due to light, preference of side and other visual effects. The tank (28 cm × 18 cm) contained 1 L of artificial sea water with pH 7.7 or 8.1. Crabs were individually caged with a plastic cylinder in the middle of the tank and filter papers (1 cm^2^) were dropped on either side of the tank. For the negative control experiment, both filter papers were blank. In the three PEA conditions, the filter paper on one side contained 200 μL of the respective PEA concentration (3 · 10^−6^ mol/L, 3 · 10^−5^ mol/L and 3 · 10^−4^ mol/L) while the paper was left empty on the other side of the tank (control). The side of the PEA filter paper was randomized. After allowing the cue to diffuse for 20 seconds, the crab was released by lifting the plastic cylinder and observed for 2 minutes. For the data analysis, the tank was virtually divided into thirds (see Fig. 3) and the time spent in each of the three area was recorded manually or by video to ensure consistency.

**Figure 3:**
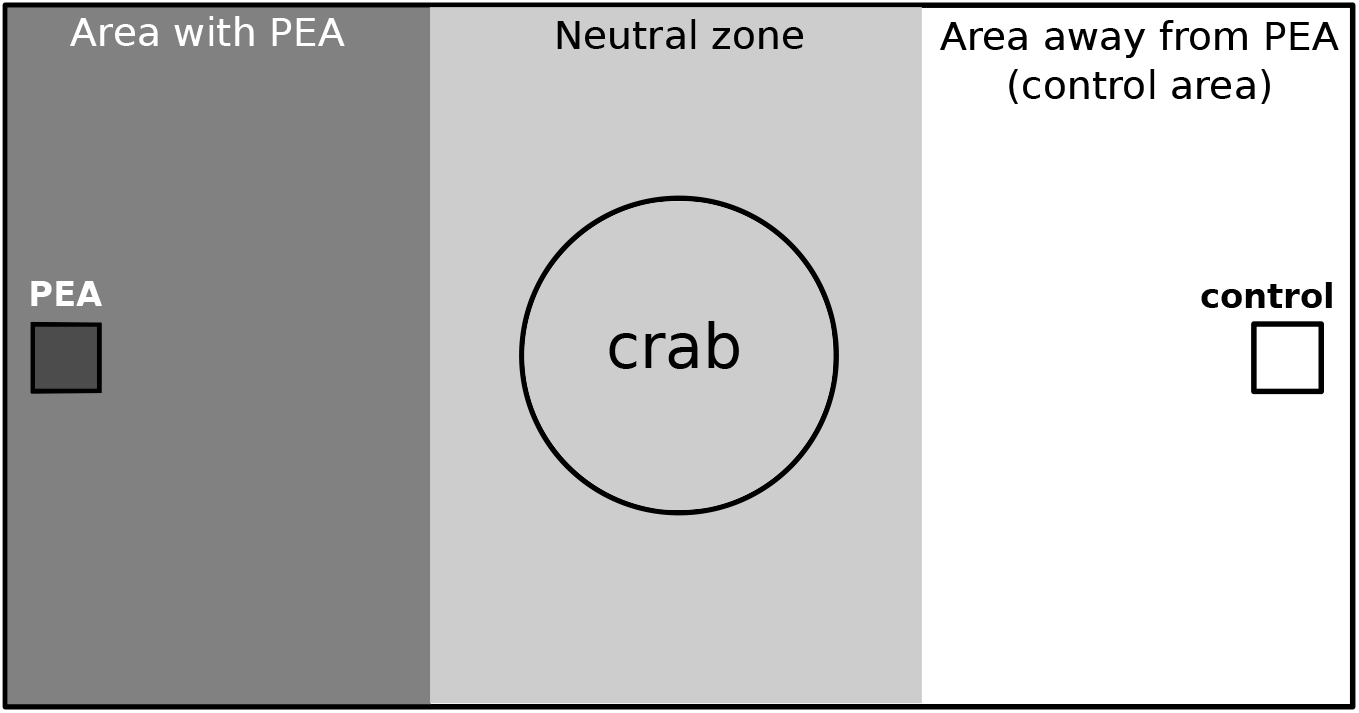
Set-up of the behaviour experiment. The hermit crab was caged with a plastic cylinder in the middle of the neutral zone whilst the cues were dropped on either side of the tank on filter paper (squares). After 20 seconds of diffusion time, the cylinder was lifted and the movement pattern of the hermit crab was observed and recorded by video for 2 minutes. For data analysis, the time spent in the three areas (dark gray, gray, white) of the tank was stopped.

By manipulating the pH of the bioassay water with HCl instead of CO_2_ we are able to focus on the effect of pH and manipulate only the proton concentration in the water (Gattuso et al. 2010). Leaving the carbon chemistry untouched allows us to study the isolated effect of pH on the behaviour of hermit crabs. Thereby we can directly compare the effect of pH on the response of hermit crabs to PEA with the changes in conformation and charge of the signalling cue. This allows us to improve our understanding of the underlying mechanisms.

### Statistics

All statistics were carried out with R (version 3.2.3-4) (R Core Team 2015). The behaviour assay as described above has a paired structure within the level of the individual crab as the same hermit crab was tested four times (negative control, three PEA concentrations). Applying a linear mixed effect model (‘lme4’ R package) (Bates et al. 2015) the effect of the concentration of PEA on the time spent in either area was tested for both pH scenarios (pH 7.7 and 8.1). Multiple comparisons of the response to PEA at different concentrations were carried out with Tukey’s Test using the ‘multcomp’ package in R (Hothorn et al. 2008).

### Computations of PEA

The conformation of neutral and protonated PEA was studied using quantum chemical methods. The calculations were carried out with the supercomputer (Viper) of the University of Hull and validated by NMR spectroscopy. Calculating the total energy of geometry-optimised PEA at different constrained torsion angles *τ* of the amino side chain allowed us to determine energetically favourable conformations and barrier heights (torsion angle energy scan). Due to the structure of PEA (Fig. 2), the position of the amino side chain is the main structural factor in the determination of energetically favoured geometries.

However, the aquatic environment in which PEA is dissolved also has to be taken into account. Long-range and short-range interactions with water can have a large impact on the conformation (Roggatz et al. 2018). Therefore, the torsion angle energy scan was studied in gas phase, in a dielectric infinite continuum of water using the CPCM approach (Barone and Cossi 1998; Cossi et al. 2003, implicit solvent approach) implemented in ORCA version 4.0.1 (Neese 2012, 2018) and in an implicit solvent environment including the interaction of one explicit water molecule with the amino group. One explicit water molecule per ionisable group was found to improve accuracy and reliability of isotropic nuclear magnetic shielding calculations (Roggatz et al. 2018). All eigenvalues of the identified minima were checked for imaginary frequencies.

Geometry optimisations in ORCA (version 4.0.1) were performed with the PBE0 exchange correlation functional (Adamo and Barone 1999) using a pc-2 basis set (Jensen 2001, 2002a,b). D3 dispersion correction (Grimme et al. 2010, 2011) was included and the RIJ-COSX approximation (Neese et al. 2009) with a def2/J auxiliary basis set (Weigend 2006) was applied. The final point energy values for each geometry optimised conformation were then plotted against the respective constrained torsion angle. To facilitate the comparison between the different environments, the total energy is shown relative to the minimum energy in the *gauche 1* conformation.

To obtain the molecular electrostatic potential (MEP) of selected conformations of PEA, the GAMESS program (Schmidt et al. 1993, version 18/08/2016, R1) was used with the PBE0 exchange functional in conjunction with the pc-2 basis set (Jensen 2001, 2002a, b). Calculations were carried out with a polarisable continuum model. Using the wxMacMolPlt program (Bode and Gordon 1998, version 7.6), a three-dimensional electron density isosurface was created with 100 grid points and a contour value of 0.1 e·a_0_^−3^. To visualise the MEP, the density isosurface was coloured with a maximum value of 0.9 E_*h*_·e^−1^ and the RGB colour scheme with red representing positive, green neutral and blue negative charge.

Isotropic nuclear magnetic shielding values of ^1^H nuclei were calculated with ORCA (version 3.0.0) at the PBE0/aug-pc-2 level of theory. As for the geometry optimisations, RIJ-COSX approximation with a def2-TZVPP/J auxiliary basis set was used. All nuclear shieldings calculations were run with the individual gauge for localized orbitals method (IGLO) (Kutzelnigg et al. 1990). The resulting nuclear shielding constants are compared to experimentally determined chemical shifts (see below).

### NMR Spectroscopy

Samples for NMR measurements were prepared with 2-phenylethylamine hydrochloride (Sigma-Aldrich, 98%) at pH 11.8 and pH 6.8 to ensure a high percentage (>99%) of PEA in the respective protonation state. PEA was measured at 0.05 mol/L in 0.04 mol/L sodium phosphate buffer to stabilise the pH during measurement with 10 % deuterium oxide (Sigma-Aldrich) as solvent lock. NMR spectra were recorded on a JEOL ECZ 400S spectrometer with Tetramethylsilane (TMS, 50μL, Sigma-Aldrich, >99.5%) *δ_H_* = 0 as the internal standard.

### Computations of Receptor Binding

32 amino acids, known to be involved in the ligand binding site (Cichero et al. 2013), were cut out of the human homology model of the TAAR1 receptor in the active state from the GPCR database (Pándy-Szekeres et al. 2018). The position and size of the chosen binding pocket is shown in Fig. 4a. Using Avogadro (Hanwell et al. 2012, version 1.1.1), peptide bonds were added to cap the cutting sites. For this, a methyl group was introduced on the amino side, a nitrogen-methyl group was added to the unbound carboxyl side and hydrogens were added to the amino acid backbone. The molecular capping procedure of amino acids was adapted from the MFCC approach (Zhang and Zhang 2003).

**Figure 4:**
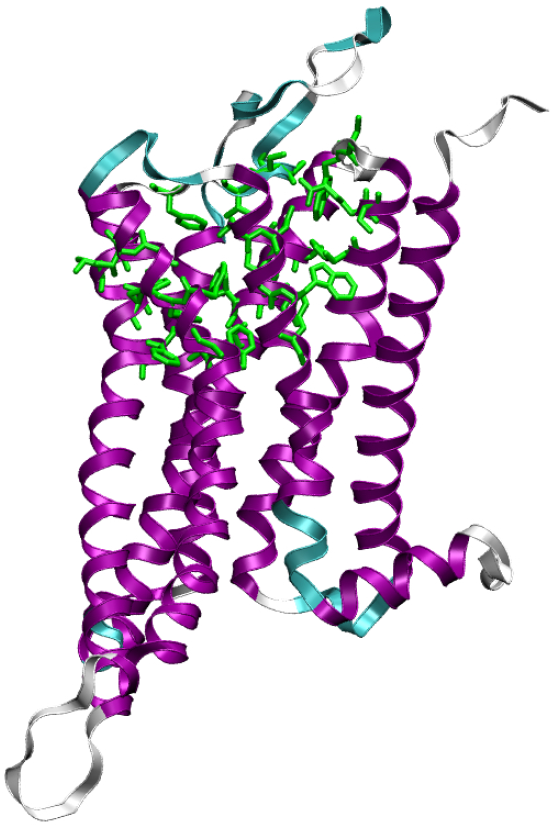
Secondary structure of the TAAR1 receptor with the alpha helix in purple, turns in cyan and coils in white. The amino acids of the binding pocket are shown in green stick representation. The majority of the chosen binding pocket is part of the alpha helix.

PEAH^+^ was positioned in the center of mass of the binding pocket in the folded conformation facing Asp103, as this was the binding point in previous studies (Cichero et al. 2013). All added methyl groups, nitrogen and hydrogen atoms were allowed to optimise using universal force field (Rappé et al. 1992) in Avogadro. Keeping PEA and the amino acid backbone constrained, the manually added atoms were then reoptimised using the PBE functional (Perdew et al. 1996) with D3 dispersion correction (Grimme et al. 2010, 2011) and SZV-MOLOPT-GTH basis set (VandeVondele and Hutter 2007; Krack 2005) in CP2K (VandeVondele et al. 2005; Hutter et al. 2014, version 6.1).

In the following, the binding pocket was placed inside a spherical cavity with radius 18.12 Å, outside which a dielectric continuum simulated the protein background. For this, the self-consistent reaction field (SCRF) method (Wong et al. 1991) with a dielectric constant of 8 was applied. Previous studies showed that the protein environment in an enzymatic reaction can be adequately represented by embedding the active site in a dielectric cavity with the dielectric constant of 8 (Siegbahn and Himo 2009). The size of the cavity for the TAAR1 receptor binding pocket was chosen to contain the van der Waals radii of all atoms. PEA was then allowed to optimise inside the constrained binding pocket using the PBE (Perdew et al. 1996) functional with D3 dispersion correction (Grimme et al. 2010, 2011) and TZV2PX-MOLOPT-GTH basis set (VandeVondele and Hutter 2007; Krack 2005) in CP2K. Thereby, the poisson solver (Blöchl 1995; Martyna and Tuckerman 1999) was used to isolate the (40 Å)^3^ boxes in the periodic environment. By default, CP2K introduces a uniform background charge in charged periodic systems.

To determine the effect of the charge of PEA on the receptor binding, PEAH^+^ and neutral PEA were optimised with the same starting conformation and position. For neutral PEA, all three positions of the electron lone pair in the starting conformation were compared and the energetically favoured conformation was chosen.

## Results

### Role of PEA for Hermit Crabs

The choice experiment with hermit crabs reveals that PEA is an attractant for hermit crabs. In pH 8.1, 12 out of 20 hermit crabs spent more time in the area with the cue (3 · 10^−4^ mol/L) than during the respective negative control experiment and 14 out of 20 hermit crabs preferred the area with PEA in pH 7.7. Fig. 5 shows that whilst the time spent in the third of the tank furthest away from the cue decreased in a clear dose-dependent manner (Fig. 5a), the time spent in the area with PEA increased subsequently (Fig. 5b).

**Figure 5:**
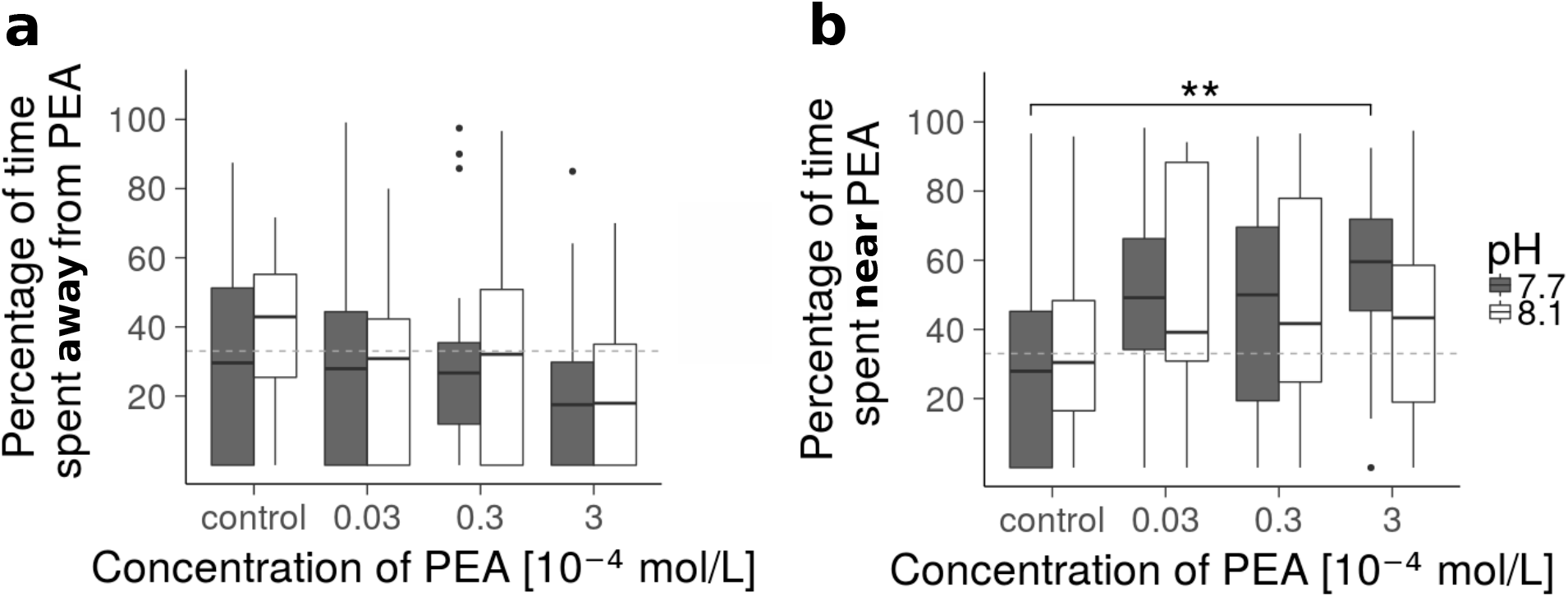
Behavioural response of hermit crabs (n=20) to PEA at current pH (8.1) and end-of-the-century level (7.7). Percentage of time spent in the third of the tank furthest away from PEA (a) and near the PEA source (b) at different PEA concentrations. The dashed gray line indicates a third of the time. At pH 7.7 (dark gray), hermit crabs spent significantly more time near the highest dose (b, *p <0.01*, indicated by bracket with asterisks). There was no significant difference at pH 8.1 (white). The boxplot depicts the median with first and third quartile of the distribution. Whiskers extend to 1.5× the interquartile range; data beyond that range are defined as outliers and plotted individually.

The response of hermit crabs in the control experiments are comparable in both pH conditions, indicating that the decreased pH in the absence of a signalling cue had no measurable effect on their behaviour. The attractive effect of PEA was stronger at low pH. At pH 7.7, the hermit crabs spent significantly more time in the area with PEA during the highest exposure experiment (3 · 10^−4^ mol/L) than during the negative control experiment (*post-hoc Tukey’s test, p<0.01*, on average 27%). On the other hand, the dose-dependent response to PEA was not significant at pH 8.1 (*chi-squared test, p=0.29*). The date of the experiment had no significant effect on the model, testifying the rigorousness of the procedure. A table of the recorded times and conditions of the behaviour experiment can be found in Online Resource 1.

### Conformation and Charge Distribution of PEA at Different pH and in Different Environments

The 360° torsion angle energy scans shown in Fig. 6 reveal three energetically favoured conformations for both, neutral and protonated PEA. An extended (*anti*) and two folded (*gauche*) conformations were found to be energetic minima in all three environments (Table 1). It is important to note that especially the *gauche 1 /gauche 2* energetic differences are very close to chemical accuracy, making a precise conformational prediction difficult.

**Figure 6:**
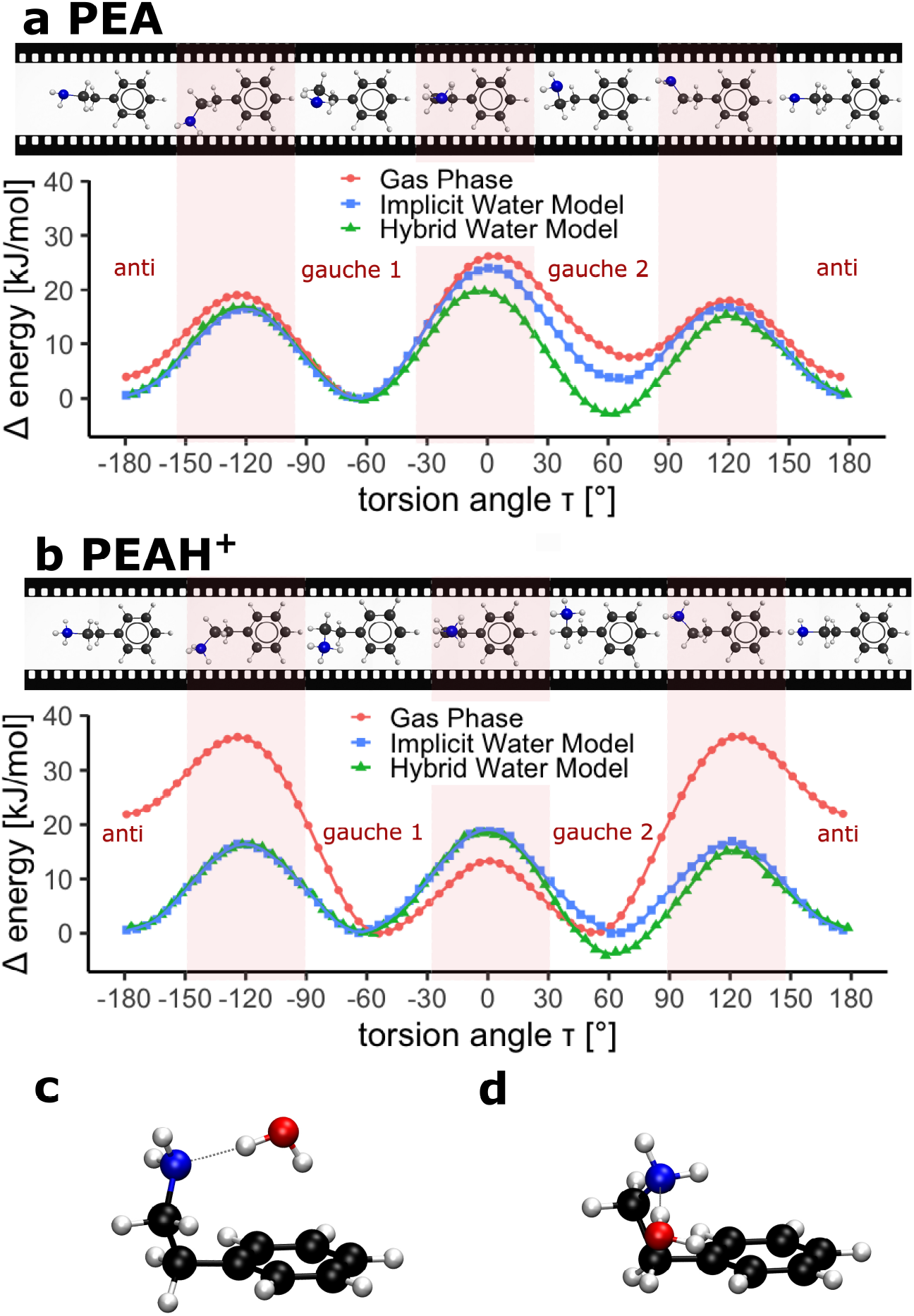
Energy scan around the amino side chain torsion angle of neutral PEA (a) and PEAH^+^ (b) and interactions of neutral PEA with water (c & d). Energy values are relative to the minimum energy in the *gauche 1* conformation. Conformations for selected torsion angles are depicted in a filmstrip above the scan. Favoured geometries are energetic minima; both protonation states show minima at an extended (*anti*) and two folded (*gauche*) conformations. The potential energy curve is shown for two solvation models and gas phase (red circles). The implicit water model (blue squares) is extended by including the interaction of one explicit water molecule with the amino group (hybrid water model, green triangles). As only one water molecule is added, this scan is asymmetric. The two *gauche* conformations in the hybrid water model of neutral PEA are compared in (c) and (d). (c) corresponds to the energetically favoured conformation *gauche 2* whilst (d) is the conformation that was the most stable in the implicit water model (*gauche 1*). Hydrogen atoms are depicted in white, carbon atoms in black, nitrogen atoms in blue and oxygen atoms in red.

**Table 1:**
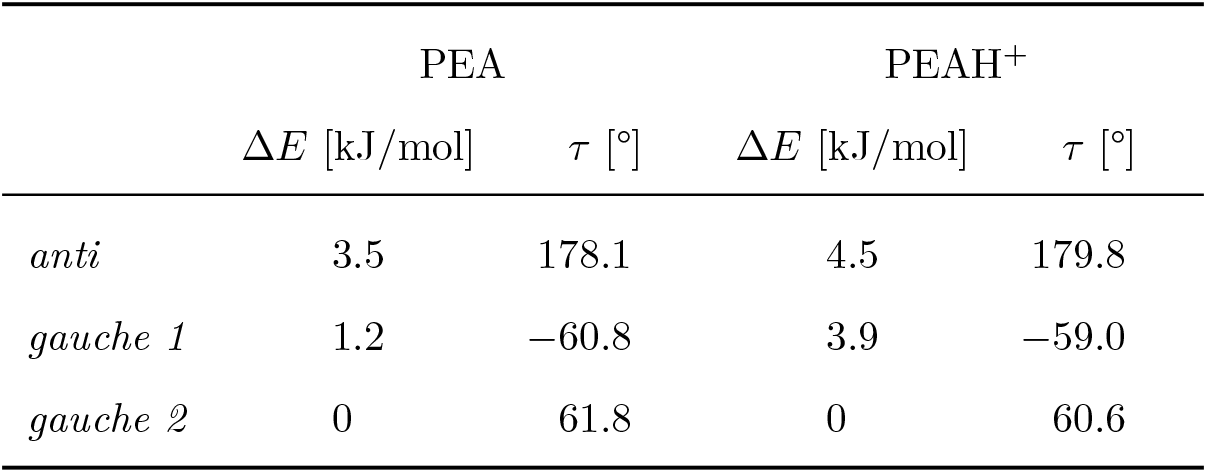
Torsion angles *τ* and energy differences Δ*E* of the energetically favoured conformations of PEA in solvation as determined with the hybrid water model. The energetic minima were optimised without geometrical constraint and the energy of the explicit water molecule was substracted. Energy differences are shown relative to the lowest energy conformation.

|  | PEA |  | PEAH<sup>+</sup> |  |
| --- | --- | --- | --- | --- |
| | $\Delta E$ [kJ/mol] | $\tau$ [°] | $\Delta E$ [kJ/mol] | $\tau$ [°] |
| *anti* | 3.5 | 178.1 | 4.5 | 179.8 |
| *gauche 1* | 1.2 | −60.8 | 3.9 | −59.0 |
| *gauche 2* | 0 | 61.8 | 0 | 60.6 |

Solvation has a clear effect on the scan profile. This results in different relative stabilities of the energetic minima with slightly differing torsion angles and different barrier heights depending on the environment. Fig. 6a & b compare the energy scans as a function of the torsion angle in three environments: in gas phase, in a conductor-like polarisable continuum model (implicit water model) and in the implicit water environment including one explicit water molecule interacting with the amino group (hybrid water model).

The neutral PEA scan (Fig. 6a) shows an asymmetry in the relative stability of the two folded conformations (*gauche 1 τ* ≈ −60°, *gauche 2 τ* ≈ +60°). This can be explained by the asymmetry of the amino group, containing two hydrogen atoms (H_N_) and an electron lone pair. In the *gauche* conformation either H_N_ or the nitrogen lone pair are interacting with the aromatic ring, leading to different relative stabilities. The environment, however, has a clear impact on these relative stabilities. In the implicit water model, the *gauche 2* conformation with lone pair-*π* interaction (*τ* ≈ +60°) is stabilised compared to the gas phase. This effect becomes more pronounced with the inclusion of an explicit water molecule (hybrid water model). This reverses the relative stabilities of the folded conformations. Whilst *gauche 1* was the stable folded conformation in gas-phase and in the implicit water model, *gauche 2* is favoured in the hybrid water model. This is due to the hydrogen atoms of the water molecule forming hydrogen bonds with the nitrogen lone pair and the *π*-system on either side and thereby promoting the energetic stability (Fig. 6c). The distance between the amino group hydrogen atom closest to the benzene ring and the *π*-system is 3.28 Å for the *gauche 1* conformation (Fig. 6d) whilst the distance between the *π*-system and the water hydrogen atom is 2.48 Å (*gauche 2*, Fig. 6c). As oxygen atoms are known to be rather weak hydrogen-bond acceptors compared to nitrogen atoms (Böhm et al. 1996), the nitrogen lone pair of PEA interacts with the water hydrogen atom rather than PEA forming a bond between the nitrogen hydrogen atom and water oxygen atom.

Overall, solvation has a strong effect on PEAH^+^ (Fig. 6b). The difference between the gas phase model and the solvation models are particularly pronounced in the relative stability of the extended conformation (*τ* ≈ ±180°) and the barrier height between the folded and extended conformation. The implicit water model stabilises the *anti* conformation. Adding one explicit water molecule (hybrid water model) decreases the relative energy of one of the folded conformations. Similarly to neutral PEA, the folded conformation with the water mediating between the amino group and the aromatic ring is the most stable form for protonated PEA in the hybrid model. In this conformation, the distance between the hydrogen atom of the water molecule and the *π*-system is 2.39 Å.

The conformations of the energetic minima deducted from the torsion angle energy scans of the hybrid water model were also optimised with released torsion angle constraint. In water, *gauche* is energetically favoured over the *anti* conformation for both protonation states. After substracting the energy of the water molecule, the torsion angles and energy differences are shown in Table 1. The xyz-files of the energetic minima in gas-phase, in the implicit and hybrid water model can be found in Online Resource 2.

The torsion angle energy scans (Fig. 6a & b) are unsymmetric, reflecting the range of energetically favoured conformations of PEA. However, neutral and protonated PEA both have two degenerate (i.e. same energy level) folded conformations as their energetic minimum. By rotation around the C-N bond, the flexible amino side chain enables the molecule to lock into the same folded conformation for negative as well as positive torsion angles.

The relative population in the different conformations is governed by the energy difference and the temperature of the system. By calculating the Boltzmann factor, the energy differences (Table 1) can be translated into relative populations. With an ocean annually-averaged temperature of 17°C for 2018 (NOAA 2019), the ratio of extended to folded conformation is 1:9 for neutral PEA and 1:13 for PEAH^+^. Decreasing the pH leads to an increase in the population of the folded conformation.

Both, neutral PEA and PEAH^+^ show charge separations in the amino group and C-H bonds (Fig. 7a & b). PEAH^+^ is overall more positively charged (red in Fig. 7a & b) than PEA, whereby the difference in the charge distribution is particularly pronounced in the amino group. The dipole moment can be used as a measure of the charge separation. Even in the same folded conformation, protonated and neutral PEA differ in magnitude and direction of the dipole moment (red arrow in Fig. 7c & d). The dipole moment for folded neutral PEA is 2.5 D whilst it is 12.6 D for PEAH^+^ in the same conformation.

**Figure 7:**
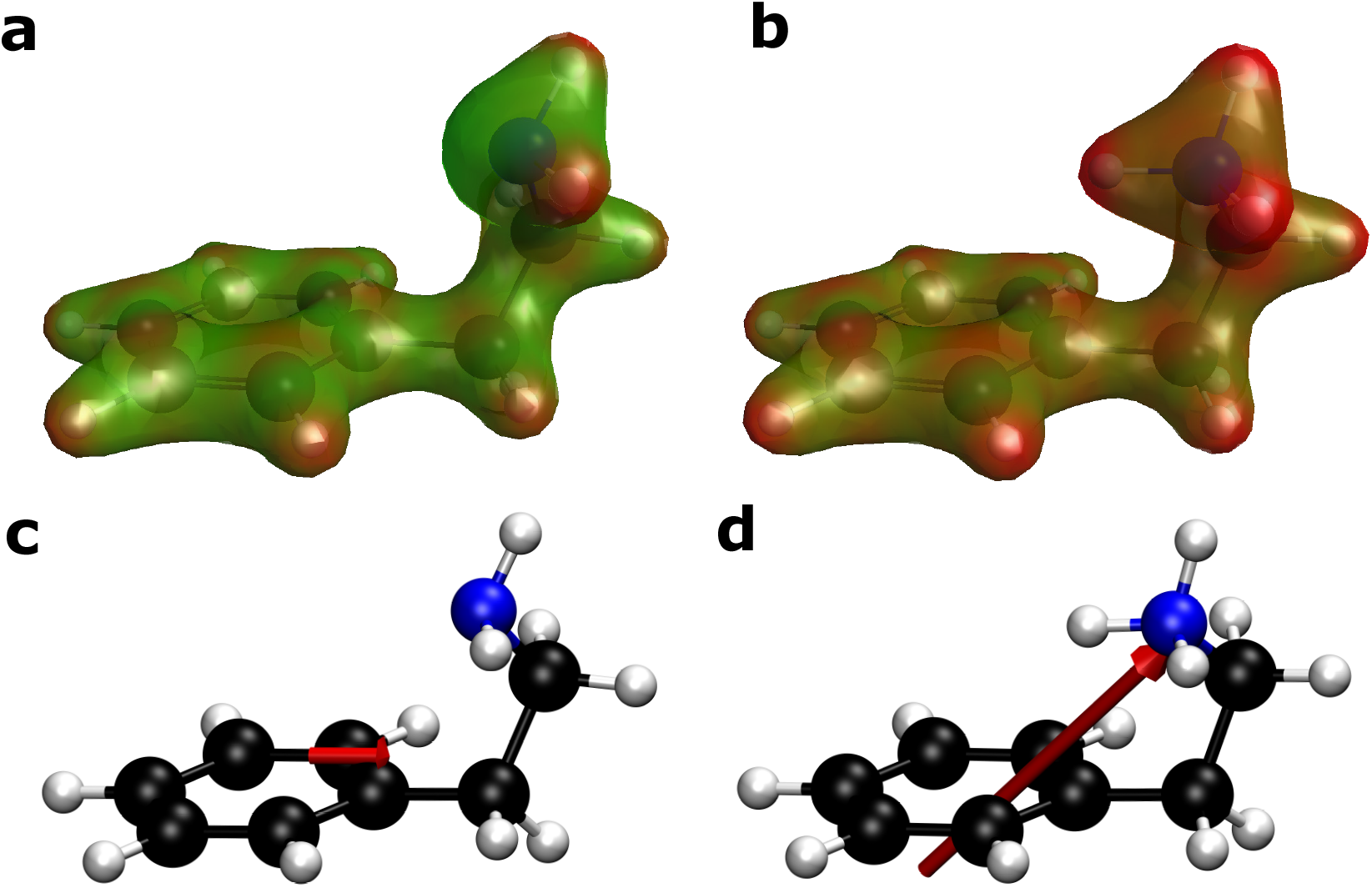
Charge distributions (a & b) and dipole moment (c & d) of folded PEA in the implicit solvation model in the protonated (b & d) and neutral state (a & c). Electron density isosurfaces (a & b) are colour-coded according to the molecular electrostatic potential with blue representing negative, green neutral and red positive charge. The 30% transparency of the electron density surface shows the conformation of the molecule underneath with hydrogen atoms in white, carbon atoms in black and nitrogen atoms in blue. The dipole moment (c & d) is represented by a red arrow pointing from negative to positive charge.

### NMR spectroscopy

To validate the quantum chemical computations of PEA, nuclear proton shieldings of the energetic minima of PEA were calculated in the hybrid water model (as described by Roggatz et al. (2018)). The calculated ^1^H shieldings were compared to experimental shifts measured in water.

The validation of the computational method using NMR spectroscopy is described further in the supplementary information. In brief, the correlation of the experimental data with the calculated shieldings for the different energetic minima were compared using linear models. This confirmed that the folded conformations, which are the computationally identified global energetic minima, fit best with the experimental values. As calculated, the folded conformation is the energetically favoured conformation in water for both protonation states. ^1^H NMR spectroscopy verifies the computational findings, showing a close fit with the hybrid water model of PEA.

### Comparison of Biological and Chemical Effect

To quantify the biological effect, the difference in the percentage of time spent near the cue at the highest concentration and the respective negative control experiment can be determined for the two pH environments. Fig. 8 shows that hermit crabs at pH 7.7 spend on average 21 % (25 ± 14 seconds) more time in the area with PEA at its highest concentration (3 · 10^−4^mol/L) than at pH 8.1 (*one-sided t-test, p=0.04*).

**Figure 8:**
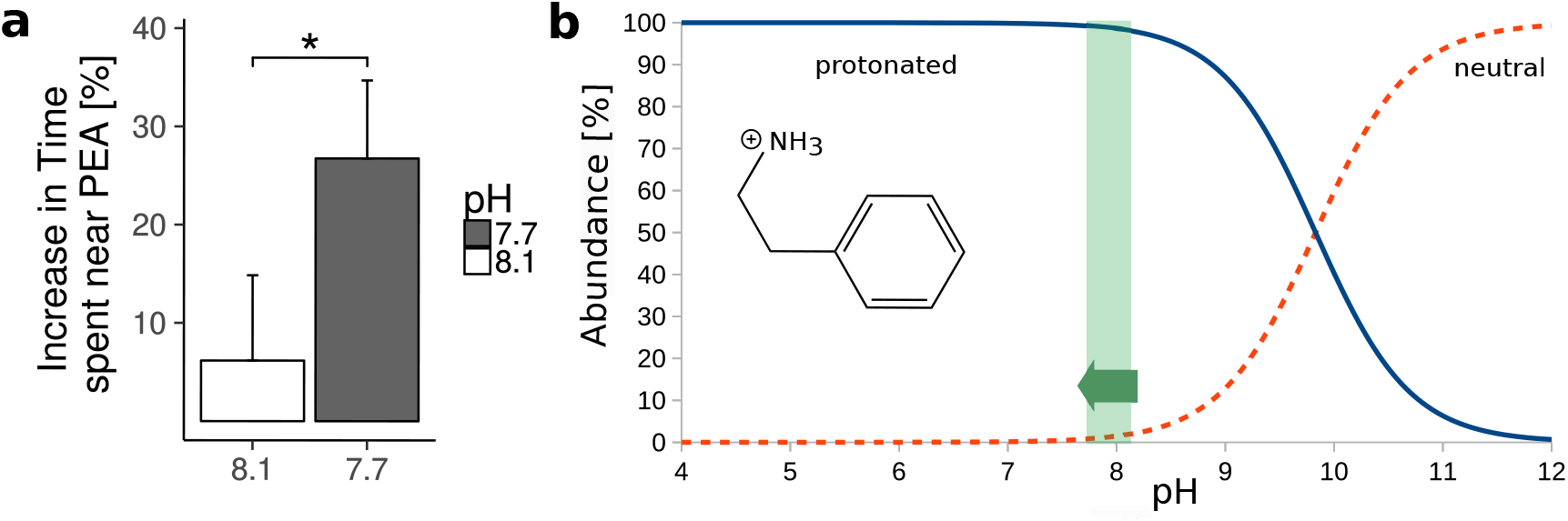
Chemical and biological effect. (a) shows the difference in time spent in the PEA area at the highest concentration and the corresponding negative control for the two pH conditions [%] with standard error bars. The asterisk indicates a significantly higher response at pH 7.7 (*one-sided t-test, p=0.04*). (b) is a plot of the Hendersson-Hasselbalch equation for PEA to visualise the proportion of the neutral (red, dashed) and protonated (blue, solid) state present across the pH range. The pH range from 8.1 to 7.7 is shaded in green.

Using the Hendersson-Hasselbalch equation (eq. 1), the percentage of protonated and neutral PEA at different pH can be estimated (Fig. 8b).

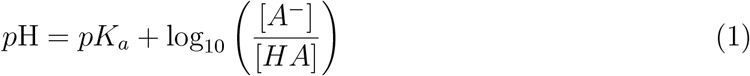

With a *pK_a_* of 9.83 (Lide 2000), PEA is mostly protonated in aquatic environments. At pH 8.1 98.0% and at pH 7.7 99.3% of PEA are in the charged state. The difference in protonation states between the two experimental conditions is therefore 1.3%.

The absolute increase in the abundance of PEAH^+^ is small. However, our calculations show that the dipole of PEA increases 5-fold upon protonation, potentially leading to an increased binding energy to its receptor.

### Receptor-Ligand Interaction in Different pH Conditions

The protonation state of PEA has a clear effect on its orientation and conformation inside the TAAR1 binding pocket (Fig 9).

**Figure 9:**
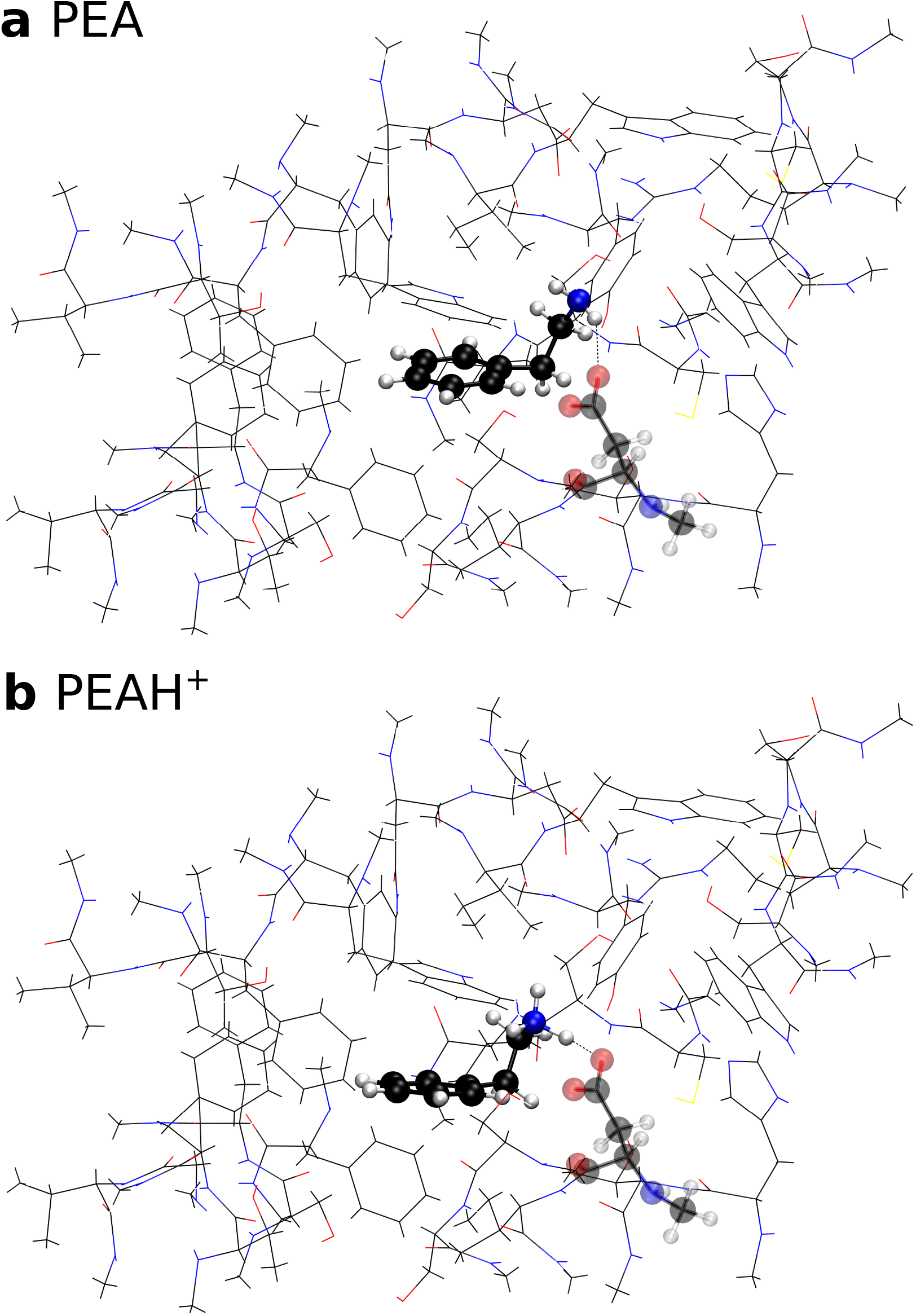
Conformation of neutral (a) and protonated PEA (b) inside the TAAR1 receptor pocket. PEA and Asp103 (transparent) are shown as ball-and-stick model, whilst the rest of the binding pocket is stylised as lines.

Starting with the geometry-optimised folded conformation of PEAH^+^ in the implicit water model at the center of mass, the root-mean-square deviation between the atomic positions of protonated and neutral PEA is 2.1 Å after geometry optimisation. The final torsion angle *τ* of the amino group side chain (see Fig. 2) of neutral PEA is 82.9° whilst PEAH^+^ adopts a torsion angle of 66.4^°^ inside the binding pocket, compared to 61.8^°^ and 60.6^°^ respectively in the non-complexed state. Furthermore, PEAH^+^ establishes a strong hydrogen bond with the negatively charged Asp103 (Fig. 9b) whilst the position of neutral PEA is mainly guided by conformational preferences (Fig. 9a). The distance between the closest amino group hydrogen atom of PEA and the negatively charged carboxyl group of Asp103 is 2.2 Å for neutral PEA whilst it is 1.4Å for PEAH^+^ (see dotted line in Fig. 9). The distance between the PEA nitrogen atom and the aspartate oxygen atom is 3.1 Å for neutral PEA and 2.5 Å for PEAH^+^. Also, the angle of the hydrogen bond between PEA and aspartate increases upon protonation from 147.2° to 169.9°. This leads to a binding energy of −548 kJ/mol for PEAH^+^ and −102 kJ/mol for neutral PEA. Hence, the binding energy of PEAH^+^ with the TAAR1 receptor site is over 5 times stronger than the respective binding with neutral PEA. The gaussian cube-files of the optimised geometries of PEA in the TAAR1 receptor pocket can be found in Online Resource 3.

## Discussion

In this study we demonstrate that PEA attracts hermit crabs, despite being a cue associated with predation for sea lampreys (Imre et al. 2014) and rodents (Ferrero et al. 2011). Furthermore, PEA was hypothesised to act as a feeding deterrent in brown and red macroalgae (Smith 1977; Van Alstyne et al. 2019). To the best of our knowledge, this study is the first to report PEA to function as an attractant.

We were also able to demonstrate that the potency of PEA as a chemical signal increases with decreasing pH. The threshold value for hermit crabs reacting to PEA at pH 7.7 lays below 3 · 10^−4^ mol/L, whilst at pH 8.1, the threshold must lay above 3 · 10^−4^ mol/L. The response of hermit crabs to PEA at the highest tested concentration was significantly higher at pH 7.7 than at pH 8.1. Although negative impacts of ocean acidification on marine animal behaviour predominate, some behaviours were shown to increase under elevated CO_2_ (reviewed in Clements and Hunt 2015). Our study indicates that PEA is one of the cues that are amplified by climate change.

Other organisms that use PEA as an actual predator cue (Imre et al. 2014), in contrast to hermit crabs, could encounter the same increased potency of the cue at decreased pH. An increasing PEA efficiency at lower pH could change predator-prey interactions under climate change. The decreased threshold response to PEA would result in prey detecting predation risks at larger distances in ocean acidification scenarios, giving predators a disadvantage, whilst enhancing the chance of survival for prey.

The biological function of this predator-prey interaction cue could depend on the inhabited ecological niche and the position in the food chain. *P. bernhardus* are scavengers and known to migrate into recently trawled areas to feed on the damaged or disturbed fauna generated by beam trawling (Ramsay et al. 1996). This suggests that predation and death associated cues are attractants for hermit crabs, which is in line with our findings, but contrasts our initial expectations of a predator-related cue response. In aquatic systems dopamine, structurally very similar to PEA, induces predator associated morphological defense in waterfleas (*Daphnia*) (Weiss et al. 2015). As this crustacean is below hermit crabs in the food chain, this suggests a relativity of chemical communication cues depending on the role of the organism in the ecological network of the habitat.

Eavesdropping on chemical alarm cues is known to occur between species that share the same predator and co-occur spatially and temporally (Mathis and Smith 1993; Anderson and Mathis 2016). However, predators can also be attracted by the chemical alarm cue of prey, the smell of injured prey (Mathis et al. 1995; Wisenden and Thiel 2002). This reverses the biological function of an alarm cue into a feeding cue depending on the species and its position in the food chain. The attraction of secondary predators by chemical alarm cues is comparable to our findings. Similarily, cadaverine and putrescine are repulsive odours for zebrafish (Hussain et al. 2013), but have been reported to be feeding cues for goldfish (Rolen et al. 2003). These examples are comparable to our results, however, to the best of our knowledge, this study is the first to suggest a predator odour to attract scavengers.

Bacteria can biosynthesise PEA from organic detrius by decarboxylating phenylalanine (Marcobal et al. 2012). This supports our current findings that PEA is an attractant for hermit crabs that are known scavengers (Nickell and Moore 1992). As bacterial degradation of bioorganic matter (such as carcasses) decreases local pH through increased CO_2_ production, the increased response of hermit crabs in low pH is plausible. Ocean acidification is known to increase bacterial degradation activity (Piontek et al. 2010). This might promote the availability and importance of PEA as a feeding cue in future oceans. The potential pathway of pH interfering with the signal source is represented by pathway 1 in our scheme (Fig. 1). This is of particular interest as the efficacy of fresh food indicators such as amino acids and peptides broadly decreases with ocean acidification (Porteus et al. 2018; Roggatz et al. 2016; de la Haye et al. 2012; Velez et al. 2019). Assuming PEA acts as a detrius cue at low pH, its increased potency could indicate that a potential shift in the diet of hermit crabs in response to ocean acidification is possible. This raises hope for adaptation processes to climate change pressures of olfactory disruption.

Coastal ecosystems experience tidal, seasonal and annual natural fluctuations in acidity regardless of climate change (Wolfe et al. 2020). However, ocean acidification is known to reduce buffering of pH cycles (Pacella et al. 2018; Kwiatkowski and Orr 2018). Coastal diel pH extremes are expected to exceed open-ocean average pH changes for the end of the century (Pacella et al. 2018). Hermit crabs are therefore already today experiencing pH conditions that amplify the attraction to PEA, with climate change however its relevance is expected to increase.

Furthermore, the amplified potency of PEA at decreased pH suggests that the protonated state is the bioactive form. This is supported by the structural similarity of PEA to neurotransmitters, which operate at pH levels around 7.4 in human blood, where, following eq. 1, 99.4% of PEA is protonated.

As recent studies show, olfactory disruption due to ocean acidification can be closely associated with structural changes of the odour molecule (Velez et al. 2019; Roggatz et al. 2016). In our scheme (Fig. 1) this is represented by the second pathway of the pH influencing the signalling transmission. As these molecules are operating in marine environments, it is crucial to advance our understanding of solvation models to enable more realistic investigations at molecular level. Research into quantum chemical methods of ecologically relevant systems is largely underrepresented. Modelling short-range and long-range interactions of water is a trade-off between computational cost and accuracy. Implicit water models are the simplest implementation of solvation effects and computationally inexpensive. However, they only model long-range interactions with water. Our results show that the gas-phase calculations deviate from the solvation models (Fig. 6, see also Roggatz et al. (2018)). We show that solvation stabilises the extended conformation of PEA. Especially for small molecules, short-range effects of explicitly included water molecules can add important structural information to the model. Thereby, our model coincides with findings of Bouchet et al. (2016), where the same distance between the water hydrogen atom and the *π*-system (2.39 Å) was identified for the energetic minimum at absolute zero. However, this neglects the entropy and other energetic contributions to the Gibbs free energy. As Bouchet et al. (2016) shows, these can influence the favoured position of water in the folded conformation and might be crucial for the correct representation of solvation effects. Nevertheless, for PEA the energy difference between the conformers (Table 1) is comparable to those at room temperature when the Gibbs free energy is considered (Bouchet et al. 2016). Further studies are being conducted to extend the current solvation models and promote our understanding of the conformation of small molecules such as odour cues in water.

The quantum chemical calculations for protonated and neutral PEA reveal differences in the conformation (Fig. 6) and the dipole moment (Fig. 7) for the different protonation states. The observed conformational changes, on the one hand, are small and might not affect receptor-ligand interactions. In analogy to dopamine, the amino group could act as the anchoring point within the receptor followed by a rapid rearrangement of the conformation (Andujar et al. 2011). Considering the small change in the amount of active compound (1.3%) within the range of ocean acidification, it is unlikely that conformational changes of PEA account for the change in hermit crab behaviour. The dipole, on the other hand, increases 5-fold upon protonation. In contrast to conformational effects, increasing the electric interactions between receptor and ligand can significantly affect their interplay and accelerate the binding (discussed further below, Radić et al. (1997); Wade et al. (1998)). In mammals, PEA is a known dopamine receptor agonist (Barroso and Rodriguez 1996), that binds to the TAAR1 receptor (Pei et al. 2016) and inhibits the vesicular monoamine transporter 2 in monoamine neurons (Wimalasena 2011). However, although PEA is a neurotransmitter in mammals, a similar function in hermit crabs remains unknown.

To provide a proof of principle, we modeled the changes in the receptor-ligand interaction upon protonation of the ligand using the human TAAR1 receptor. To the best of our knowledge, this study is the first to include the effects of ocean acidification on receptor-ligand binding. Our results coincide with findings of Cichero et al. (2013), where the folded conformation of PEAH^+^ was found to bind to Asp103. Furthermore, protonated dopamine and its D2 binding pocket (Andujar et al. 2011) are similar in structure and binding mechanism to the PEA-TAAR1 complex, which allows us to compare the models. Whilst we report a folded conformation of PEA (torsion angle *τ* = 83° for neutral and 66° for PEAH^+^) when binding to Asp103 in TAAR1, dopamine also adopts a folded conformation (*τ* = 78°) in interaction with Asp86 of the dopamine D2 receptor at the global minimum (Andujar et al. 2011). Additionally, the distance between the nitrogen atom of PEA and the aspartate oxygen atom is 3.1 Å for neutral and 2.5 Å for protonated PEA, whilst similarly, the aspartate binding point of the dopamine D2 receptor establishes a distance of 2.9 Å with the nitrogen atom of dopamine (Andujar et al. 2011). Furthermore, these models of G protein-coupled receptors are within the range of heavy atom distances of the N-H…O bonds generally observed in protein-ligand interactions (de Freitas and Schapira 2017). Also, hydrogen bond angles in protein-ligand interactions were reported to peak at 130-180^°^(de Freitas and Schapira 2017), which includes our reported angles of 147^°^ for neutral and 170^°^ for protonated PEA. The quantum chemical calculations show that the 5-fold increase of the dipole upon protonation of the ligand leads to a more than 5-fold increase in the binding affinity between receptor and ligand (Fig. 9). This suggests an increased retention time at decreased pH. However, the relationship between binding affinity and retention time is not necessarily linear. The strength of the receptor-ligand binding affects dissociation and association constants differently (Pan et al. 2013). Radić et al. (1997) demonstrated the importance of electrostatic interactions for protein-ligand interactions by comparing the association and dissociation rates of acetylcholinesterase inhibitors. Binding with the positively charged inhibitor *m*-trimethylammoniotrifluoroacetopherone was 400-fold faster and unbinding 10-fold slower than with a neutral analogue, where a positively charged nitrogen atom was exchanged for a carbon atom in the trimethylammonium group (Radić et al. 1997).

Whilst we show that the changes in the charge of the odour molecule could account for the altered hermit crab behaviour at decreased pH, other factors also have to be taken into consideration. As shown by pathway 4 in our scheme (Fig. 1), the pH can affect the signal transduction. Ocean acidification is known to alter brain ion gradients in fish, affecting the GABA signalling pathway (Heuer et al. 2016; Nilsson et al. 2012; Williams et al. 2019). However, de la Haye et al. (2012) showed that, unlike for fish, hermit crab heamolymph showed no change in ionic concentrations that correlated with the locomotory activity when exposed to low pH conditions. This suggests an impairment of the chemoreception (represented by pathway 3 in Fig. 1) rather than an interference with signal transduction in hermit crabs (arrow 4 in Fig. 1).

This study shows that for PEA, changes in the binding affinity to the receptor could be responsible for the observed change in behaviour. This is in line with a previous study by Porteus et al. (2018) showing that ocean acidification can impair the olfactory system of marine fish. However, whether the altered chemoreception is primarily attributed to changes in the odorant molecule, the olfactory receptor structure or the olfactory epithelium, might depend on the studied system (Velez et al. 2019).

Furthermore, the basic building blocks of receptors are amino acids which are known to be sensitive to pH (Tierney and Atema 1988). As olfactory receptors are in almost direct contact with the environment, their sensitivity to external conditions seems plausible. In addition to pH related changes in receptor-ligand interactions, changes in the receptors themselves are possible. Pharmaceutical studies on G protein-coupled receptors demonstrate the effect of altered pH on receptor functioning (D’Souza and Strange 1995; Gillard and Chatelain 2006; King et al. 1997). They show that changes in just a few amino acids, caused by a drop in pH, can lead to fundamental changes in the receptor, altering association and dissociation rates of ligands. Studies on pheromone binding mechanisms in insects have also shown that pH-induced conformational changes can play a major role in the ligand-protein interaction (Di Luccio et al. 2013; Yin et al. 2015).

As the PEA receptor in hermit crabs is unknown, we assume an aspartate-based binding mechanism for our model, similar to the human TAAR1 receptor. It is important to note that solvation effects have been neglected in the receptor model. The effect of water on the binding mechanism remains to be explored. Although we don’t know the binding mechanism of PEA in hermit crabs, the demonstrated preferential binding of PEAH^+^ over neutral PEA provides an explanation of the observed behavioural change. All other potential mechanistic approaches were unable to account for the altered behaviour at increased pH. Hence, the presented model of receptor-ligand interactions is a proof of concept model that points towards a new mechanistic approach. Our current findings highlight that the change in the efficacy of a signalling cue is not necessarily linear to its protonation state abundance. Protonation can lead to changes in the dipole moment of a chemical cue, which can substantially alter protein-ligand electrostatic interactions. Subsequent changes in the receptor retention time can largely deviate in scale from the difference in protonation state abundance.

In contrast to previous studies (Roggatz et al. 2016; Porteus et al. 2018; Velez et al. 2019), we chose to manipulate only the proton concentration in the sea water. Thereby, we were able to show the isolated effect of pH-induced conformational changes on the chemical cue and its interaction with a receptor. Our findings promote the understanding of the underlying mechanisms of ocean acidification effects on chemoreception by disentangling the effects of decreasing pH and increasing CO_2_. However, increasing CO_2_ levels can have wide ranging physiological effects that are not included in this study. Building on this work, a comparison of the effect of changing pH through CO_2_ and acids could further advance our understanding of how ocean acidification interferes with the sense of smell.

## Conclusion

We were able to demonstrate that PEA is an attractant for hermit crabs whilst being widely associated as a predator cue in other animals. In addition, the response to PEA depends on the pH within a range relevant for ocean acidification scenarios by 2100. Interestingly, decreasing the pH amplifies the effect of PEA. Further research is needed to promote understanding of the role of PEA for other organisms in current and future oceans. Using quantum chemical methods we explain the effect with the changing dipole of the cue. We show that the altered electronic properties can impact receptor-ligand affinity and thereby affect the retention time of the ligand. Future research should consider modelling pH-dependent changes in the receptor site, the chemical cue as well as their interaction. This study provides a rare example of a chemically-mediated behaviour that is enhanced in future ocean conditions and showcases the power of cross-disciplinary research to help unravel the underlying mechanisms.

## Supporting information

Supplemental Resource 3

Supplemental Resource 2

Supplemental Resource 1

## Acknowledgements

We acknowledge the Viper High Performance Computing facility of the University of Hull and its support team. The authors would like to thank Dr. R. Terschak and V. Swetez for technical and animal husbandry support and Dr. R. Lewis for advice and help with NMR spectroscopy. PS is financially supported by the University of Hull, UK, as part of a PhD project cluster. CCR is supported by the ERC-2016-COG GEOSTICK project grant of Prof. D. Parsons.

## Notes

### Competing Interest Statement

The authors have declared no competing interest.

