## Supplementary for "Ocean Acidification Amplifies the Olfactory Response to 2-Phenylethylamine: Altered Cue Reception as a Mechanistic Pathway?"

### Behaviour Data

A table of the recorded times and conditions of the behaviour experiment with hermit crabs can be found in resource 1

### Optimised Geometries of 2-Phenylethylamine

The xzy-files of the optimised geometries of 2-phenylethylamine in gas-phase, implicit and hybrid solvation environment can be found in resource 2.

### Optimised Geometries of 2-Phenylethylamine in TAAR1

The gaussian cube-files of the optimised geometries of 2-phenylethylamine in the TAAR1 receptor pocket can be found in resource 3.

### Computational Method Validation

To validate the quantum chemical computations, nuclear proton shieldings of the energetic minima of 2-phenylethylamine (PEA) were calculated in the hybrid water model (as described by Roggatz et al. (2018)). The calculated $^{1}H$ shieldings were compared to experimental shifts measured in water by plotting calculated values against experimental results and performing a least-squares linear regressions for the different conformations *gauche* and *anti* of PEA (Fig. 1).

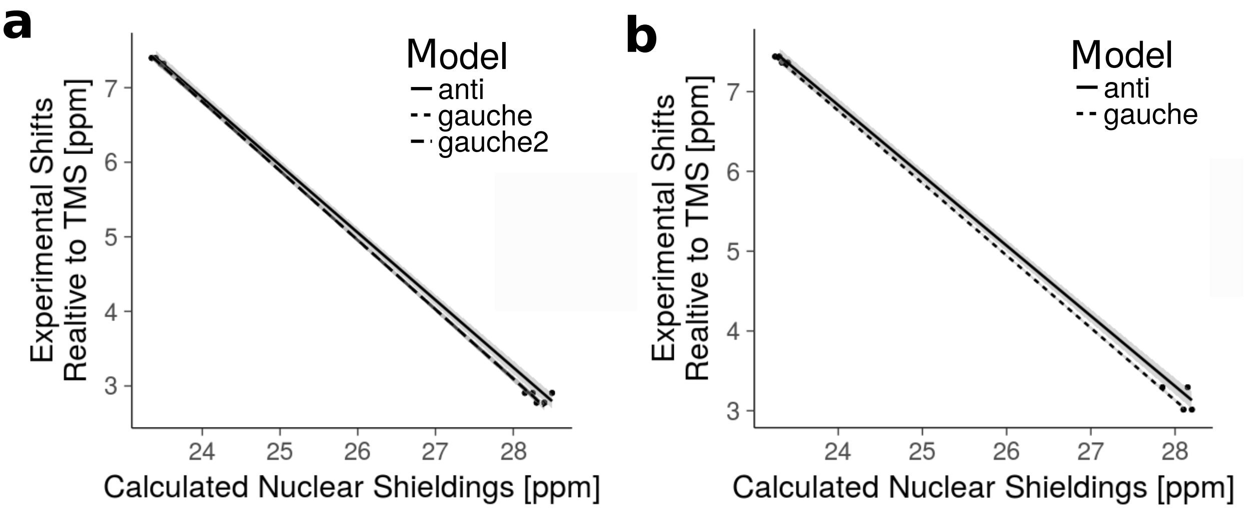


Figure 1: Calculated nuclear proton shieldings as a function of the experimental NMR $^{1}$H shifts for neutral PEA (a) and PEAH$^{+}$ (b). The 95 % confidence interval is shown in gray. Only in the protonated state, the amino protons were measurable and thus excluded from the fit.

Thereby the calculated nuclear shieldings are absolute values whilst the experimental shifts are measured relative to the pH insensitive reference TMS, for which the $^{1}$H shift is set to zero. Due to the symmetry of the ring and the free rotation about single bonds (fast exchange rate) only 4 carbon-bound $^{1}$H shift can be measured (Fig. 2). The corresponding calculated shieldings were averaged to facilitate comparability. As the protons bound to nitrogen were only observed in the low pH NMR experiments (PEAH$^{+}$) they were excluded from the linear model. $H_N$ are known to be problematic in NMR shielding calculations as they are highly affected by hydrogen bridge networks (Frank et al. 2011).

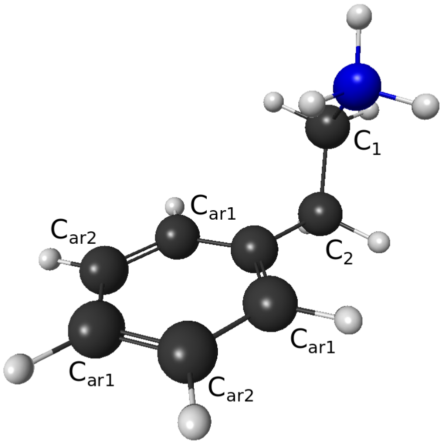


Figure 2: Structure and naming of the carbons in PEA to assign experimental $^1$H NMR shifts to the calculated shieldings for both protonation states.

With the linear regression functions (eq. 1, plotted in Fig. 1) the calculated nuclear magnetic $^1$H shieldings ($\sigma$) can be translated into chemical shifts ($\delta$) . The comparison of experimental $^1$H shifts and calculated $^1$H shifts is shown in Table 1. It becomes apparent that the difference between folded (*gauche*) and extended (*anti*) conformation of PEA are reflected mainly in the chemical proton shifts of the side chain ($C_1$-H and $C_2$-H in Fig. 2) and values from the computational models of folded PEA seem closer to the experimental results.

$$\delta = a \cdot \sigma + b \tag{1}$$

$$\begin{aligned} \text{for PEA}: \quad a_{anti} &= -0.90, & b_{anti} &= 28.55\,\text{ppm} \\ a_{gauche} &= -0.93, & b_{gauche} &= 29.12\,\text{ppm} \\ a_{gauche2} &= -0.93, & b_{gauche2} &= 29.10\,\text{ppm} \\ \text{for PEAH}^+: \quad a_{anti} &= -0.88, & b_{anti} &= 27.96\,\text{ppm} \\ a_{gauche} &= -0.91, & b_{gauche} &= 28.53\,\text{ppm} \end{aligned}$$

Table 1: Comparison of experimental $^1$H shifts with computational shifts of protonated and neutral PEA in water. The naming of the carbons of PEA is visualised in Fig. 2.

| | neutral PEA [ppm] | | | | PEAH$^+$ [ppm] | | |
|---|---|---|---|---|---|---|---|
| | experimental | calculated | | | experimental | calculated | |
| | | *anti* | *gauche* | *gauche 2* | | *anti* | *gauche* |
| $C_1$-H | 2.91 | 2.80 | 2.86 | 2.96 | 3.30 | 3.18 | 3.27 |
| $C_2$-H | 2.78 | 2.89 | 2.82 | 2.72 | 3.02 | 3.13 | 3.04 |
| $C_{ar1}$-H | 7.32 | 7.32 | 7.31 | 7.31 | 7.37 | 7.36 | 7.37 |
| $C_{ar2}$-H | 7.40 | 7.41 | 7.42 | 7.42 | 7.44 | 7.45 | 7.44 |
| TMS-H | 0 | 0.27 | 0.03 | 0.03 | 0 | 0.40 | 0.14 |

The two folded conformations (*gauche*) of neutral PEA are not separable in the linear regressions (see Fig. 1). However, the folded and extended conformation show different linear fits for both protonation states. To measure the correlation between experimental NMR shifts and calculated shieldings, the accuracy of the fit can be determined. As the calculated values are absolute but the experimental shifts are measured relative to TMS, the calculated $^1$H NMR shielding for TMS corresponds to the x-axis intercept. The accuracy of the fit can be estimated on the basis of the extrapolated experimental value for the calculated $^1$H TMS shielding (Roggatz et al. 2018) (see last row in Table 1). Thereby, a perfect model fit would have the accuracy 0 ppm: The calculated TMS proton shielding (31.3 ppm) would coincide with the experimental NMR measurement where TMS was used as a reference (i.e. set to zero). Estimating the accuracy of the fit of the experimental findings with the computational models allows us to validate the quantum chemical calculations.

With an accuracy of ±0.03 ppm, the *gauche* conformations of neutral PEA fit the experimental values better than the *anti* conformation (accuracy ±0.27 ppm). Likewise in the protonated state, the *gauche* conformation of PEAH$^+$ reaches an accuracy of ±0.14 ppm while the accuracy of the computational model of PEAH$^+$ in the *anti* conformation is only at ±0.40 ppm. For both protonation states, the folded conformation fits the experimental findings best.

Additionally, the scattering of values in the linear model can give valuable information on the goodness of fit. The 95 % confidence intervals (gray band in Fig. 1) of the linear models of the folded conformations (*gauche*) are much narrower for both protonation states, indicating a better fit of this computational model with the experimental results. For $PEAH^+$ the linear model of the *gauche* conformation reaches a coefficient of determination of $R^2$=99.99 % for protonated and $R^2$=99.97 % for neutral PEA, whilst the *anti* model only resulted in $R^2$=99.86 % for protonated and $R^2$=99.89 % for neutral PEA. PEA in the *gauche* conformation shows the best fit with the experimental data. This coincides with the conclusion based on the accuracy of the models (see above). The confidence intervals of the models are separable towards higher calculated shieldings. This is important to note as the accuracy of the fit was measured at 31.3 ppm (calculated $^1H$ TMS shielding), where the extrapolated confidence intervals are clearly separable.

To conclude, $^1H$ NMR spectroscopy verifies the computational findings. A comparison of the accuracies as well as the linear model fits confirm that the identified global energetic minima fit best with the experimental results. The folded conformation of PEA is the energetically favoured conformation in water in both protonation states.
